# Emodin: an alveolar macrophage protector in acute pancreatitis induced lung injury

**DOI:** 10.1101/2024.03.12.584699

**Authors:** Zhe Chen, Xuanchi Dong, Yongwei Song, Bowen Lan, Yalan Luo, Haiyun Wen, Hailong Li, Hailong Chen

**Affiliations:** Department of General Surgery, The First Affiliated Hospital of Dalian Medical University, Dalian, China; Department of General Surgery, Third People’s Hospital of Dalian, Dalian Medical University, Dalian, China; Laboratory of Integrative Medicine, The First Affiliated Hospital of Dalian Medical University, Dalian, China; Institute (College) of Integrative Medicine, Dalian Medical University, Dalian, China; Department of Psychology, University of South Carolina, Columbia, SC USA

**Author notes:** Correspondence: Hailong Li, Hailong Chen. These authors contributed equally to this work (Zhe Chen, Xuanchi Dong).

**Keywords:** Emodin, Alveolar macrophage, Mitophagy, Pyroptosis, Acute pancreatitis, Lung injury

## Abstract

Emodin (EMO), an anthraquinone derivative from roots and leaves of various plants, has been widely used in many inflammatory diseases. Alveolar macrophages (AMs) play a critical role in maintaining alveolar homeostasis in the lung. To investigate the pathophysiological mechanism of AMs in acute pancreatitis-associated lung injury (AP-ALI) and the potential protective therapeutic of EMO for AP-ALI, AMs isolated from AP-ALI mice, murine cell line MH-S and RAW264.7 were pre-treated with EMO to assess its protective roles on regulating inflammation, pyroptosis, and mitophagy of macrophages in AP-ALI. The results showed that 1) in vivo, the relative quantity of AMs was significantly decreased across the time in API-ALI; however, the mitochondria flux presented earlier changes than the resident AMs alteration in our experimental system. EMO pretreatment significantly alleviated the severity of lung injury and improved the damaged alveolar structure, and further reversed the reduction of mitochondria impairment in AMs in mice. 2) In vitro, EMO significantly suppressed LPS/ATP associated NLRP3 inflammasome activation, enhanced mitophagy, and protected mitochondria damage. Furthermore, the data of mitophagy inhibition by 3-MA demonstrated that EMO protective effects were partially via manipulating mitophagy-mitochondria-alveolar macrophage axis. Our data shed light on the comprehensive understanding on EMO therapeutics in AP-ALI.

## Introduction

Acute pancreatitis (AP) is a frequent reason for emergency department visits^1^. While mild to moderate forms of AP is associated with self-limiting systemic inflammatory responses in over two-thirds of cases, severe acute pancreatitis (SAP) continues to be a critical illness that requires protracted hospitalization, prolonged time spent in intensive care, and a sluggish recovery process^2,3^. AP is associated with systemic complications even persistent (multiple) organ failure. Acutally, acinar cell death, infectious inflammation, Intestinal mucosal barrier injury, as well as the cascade of inflammation could be the underlying pathophysiological mechanisms of AP^4^. Acute lung injury (ALI)/acute respiratory distress syndrome (ARDS) are the most involved complication^5^, furthermore uncontrolled inflammation of the lungs is commonly regarded as the primary etiology of ALI/ARDS^6^.

Autophagy is a complex process of intracellular degradation of senescent or mal-functioning organelles. Selective autophagy of mitochondria, known as mitophagy, is critical for eliminating damaged mitochondria^7^ and preserving the integrity of the mitochondrial network^8^. Because the energic production in mitochondrial is mainly oxygen-dependence, the capacity of mitochondrial of generating ATP will be significantly impacted under such conditions, sequentially leading to cellular dysfunction^9^. Then, dysfunctional mitochondria triggers the severe inflammatory responses which exacerbate lung injury and multi-organ failure^10^. Alveolar macrophages (AMs), a predominant immune cell localized in alveoli in humans and rodents, serve as sentinels in the lung alveolar during homeostasis. AMs can either amplify inflammation or isolate contamination^11^. Thus, we hypothesize that mitochondrial dysfunction or damages in AMs could be one of the predominant factor during the pathophysiological progress of AP-ALI.

Emodin (EMO) is a natural product that belongs to the anthraquinone derivative family. Recent studies suggest that EMO exhibits an extensive range of pharmacological characteristics, encompassing anti-cancer, hepatoprotective, anti-inflammatory, antioxidant, and antimicrobial effects^12,13^. Previously, it has been demonstrated that EMO exhibits anti-inflammatory properties^14^, specifically via regulating NLRP3 inflammasome activation^15^. However, how EMO impacts mitophagy or regulate the correlation of mitochondrial dysfunction and NLRP3 inflammasome activation during the AP-ALI is still poorly understood.

In the present study, we utilized a caerulein/LPS induced AP-ALI mouse model to investigate how EMO influences alveolar macrophages distribution and mitochondrial damage in AMs. Meanwhile, we also use in vitro macrophage models (MH-S and RAW264.7 macrophages) to explore the protective effects of EMO on pyroptosis, inflammasome activation and mitochondria damage. Furthermore, we used a mitophagy inhibitor to validate if EMO’s protective effects potentially via enhancing mitophagy activation. The purpose of the current study is to obtain comprehensive understanding about EMO’s therapeutic on maintaining AMs homeostasis in AP-ALI.

## Results

### EMO mitigates caerulein/LPS-induced AP-ALI in mice

To investigate how EMO protects and impedes the progress of AP-ALI, we generated a mouse model of acute pancreatitis indued lung injury AP-ALI by using a combined injection of caerulein and LPS. The data in Fig. 2a and b showed that EMO significantly decreased the tissue damage in lung by the assessment of histopathological staining: more intact alveolar structure, clear interspace, less inflammatory cell infiltration, and thinner interalveolar spectrum, both at 2h and 24h after the event. Furthermore, EMO also presented significant therapeutics on the wet/dry ratio of lung tissue (an index of vascular permeability to indicate the severity of pancreatitis, Fig. 2c and d).

Additionally, to investigate how EMO impacts the inflammasome activation and leads to alleviated inflammation in acute pancreatitis-associated acute lung injury, we assessed the levels of IL-1β and TNF-α in both serum and BALF, and the data in Fig. 2g-j suggested EMO dramatically decreased IL-1β and TNF-α secretion, alleviated NLRP3 inflammasome activation, altered the protein and cholesterol levels. Our results supported that EMO explored protective effects in in acute pancreatitis-induced lung injury.

### EMO reversed the reduction of mitochondria impairment in AMs in AP-ALI mice

Next, we explored the potential role of EMO on impacting native AMs during the AP-ALI. The successful induction of AP was proved above as histopathological assessment (Figs. 1a), lipase levels (Figs. 1b and c) at different time points (Figs. 1a-c). Importantly, here it was demonstrated that EMO treatment or Carbonyl Cyanide m-Chlorophenyl Hydrazone (CCCP, a mitochondrial depolarizing chemical used as a positive control) does not reverse the establish of the AP-ALI model in mice. It was well documented that AMs are abundant in the alveolar compartment of naive mice, residing exclusively on the alveolar side of the lung; the frequency of other immune cells was below 2%. First, AMs were isolated from BALF and subjected to a 10 min incubation with MitoTracker Green FM. Flow cytometry analysis was subsequently conducted (Figs. 1d). The data showed a reduction in mitochondrial density of AMs at 2 h and 24 h after the event, compared to the control group. Interestingly, administration of EMO reversed the changes of acute pancreatitis-associated mitochondrial density (Fig. 3b-d). Next, AMs were in vivo labeled using PKH26PCL fluorescent cell linker system based on its phagocytic capabilities. In the present experimental model, in BALF, over 97% of PKH26-labeled cells were AMs with a 90% efficiency rate. As ∼60% of alveolar spaces were devoid of AMs in naive mice^14^, we measured the levels of AM/alveolar among four groups. The data showed that there is no significant difference of at 2 h after LPS injection with or without EMO treatment. However, at 24 h after LPS injection, only fewer PKH26+ cells were observed in the pancreatitis group, compared with the control. Interestingly, the decrease of PKH26+ cells in pancreatitis mice was reversed by CCCP and EMO (Fig. 3e and f).

**Figure 1.**
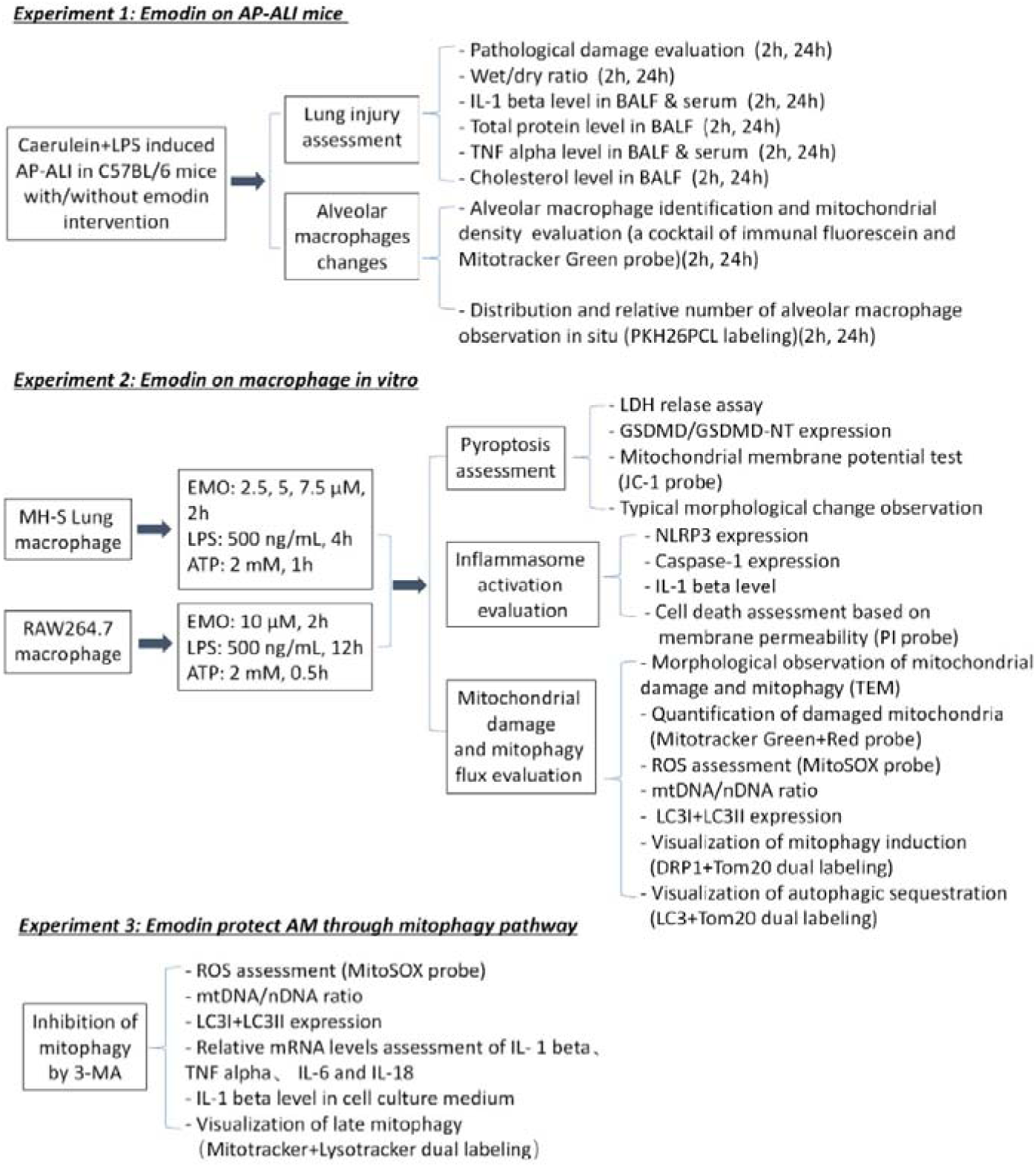
A schematic of experimental design.

### EMO inhibited the pyroptosis and mitochondrial membrane potential of macrophages in vitro

Furthermore, we extended our studies on two macrophage cell lines: MH-S lung macro-phage and RAW264.7 macrophage cells. We evaluated the cytotoxicity of EMO in vitro using the CCK-8 assay, in which vary concentration of EMO were added into MH-S and RAW264.7 cells for either 6h or 12h. The data showed that, compared to the control group, EMO present no influence on the viability of MH-S and RAW264.7 cells at dose of 20 μM or 40 μM (Figs. 2a and b). Of note, pyroptosis is recognized as a distinct process of cell death triggered by gasdermin D. This process creates holes in the plasma membrane, sequentially leads to the enlargement and membrane ballooning of cells. In the current study, EMO significantly reduced the frequency of cell swelling and membrane ballooning after 2h treatment (Fig. 4a). Furthermore, a decreased level of LDH in the culture supernatants (Fig. 4b-c), indicated the protective effect of EMO on preventing the formation of plasma membrane holes. EMO also reduced GSDMD-NT with a dose-dependent manner in MH-S cells. The results were also validated in RAW264.7 macrophage cell line (Fig. 4d-e).

**Figure 2.**
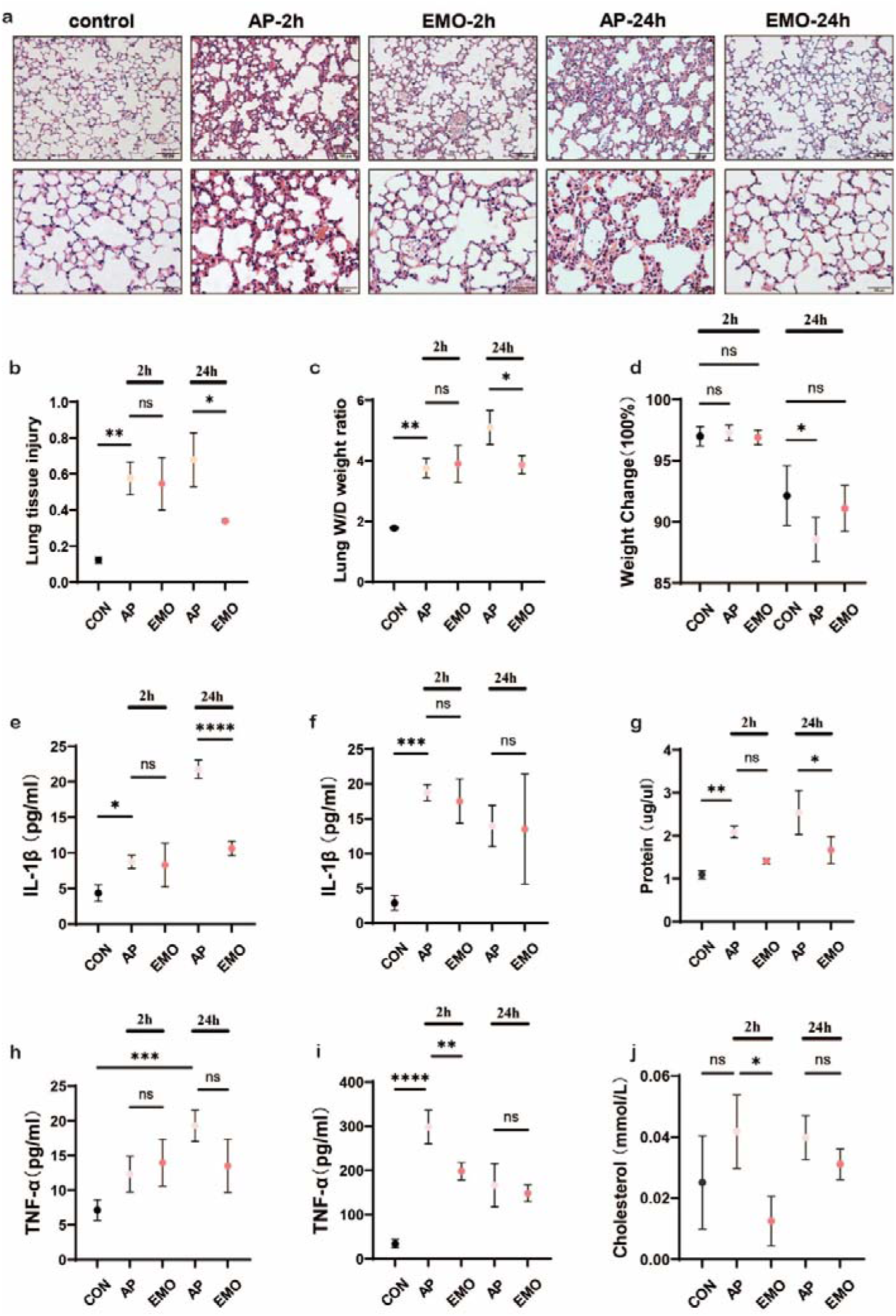
EMO mitigated caerulein/LPS-induced AP-ALI in mice. (a) Representative images of H&E staining in lung with or without EMO treatment in AP-ALI mice. (b) Quantification of lung tissue injury. n=4/group. (c) Analyze of Lung W/D weight ratio. n=4/group. (d) Measurement of body weight with or without EMO treatment. n=5/group. Expression levels of IL-1β in BALF (e) and serum (f) were evaluated by ELISA. n=4/group. (g) Protein level assessment in BALF. n=4/group. Expression levels of TNF-α in BALF (h) and serum (i) were evaluated by ELISA. n=4/group. (j) Measurement of cholesterol level in BALF. n=3∼4/group.

**Figure 3.**
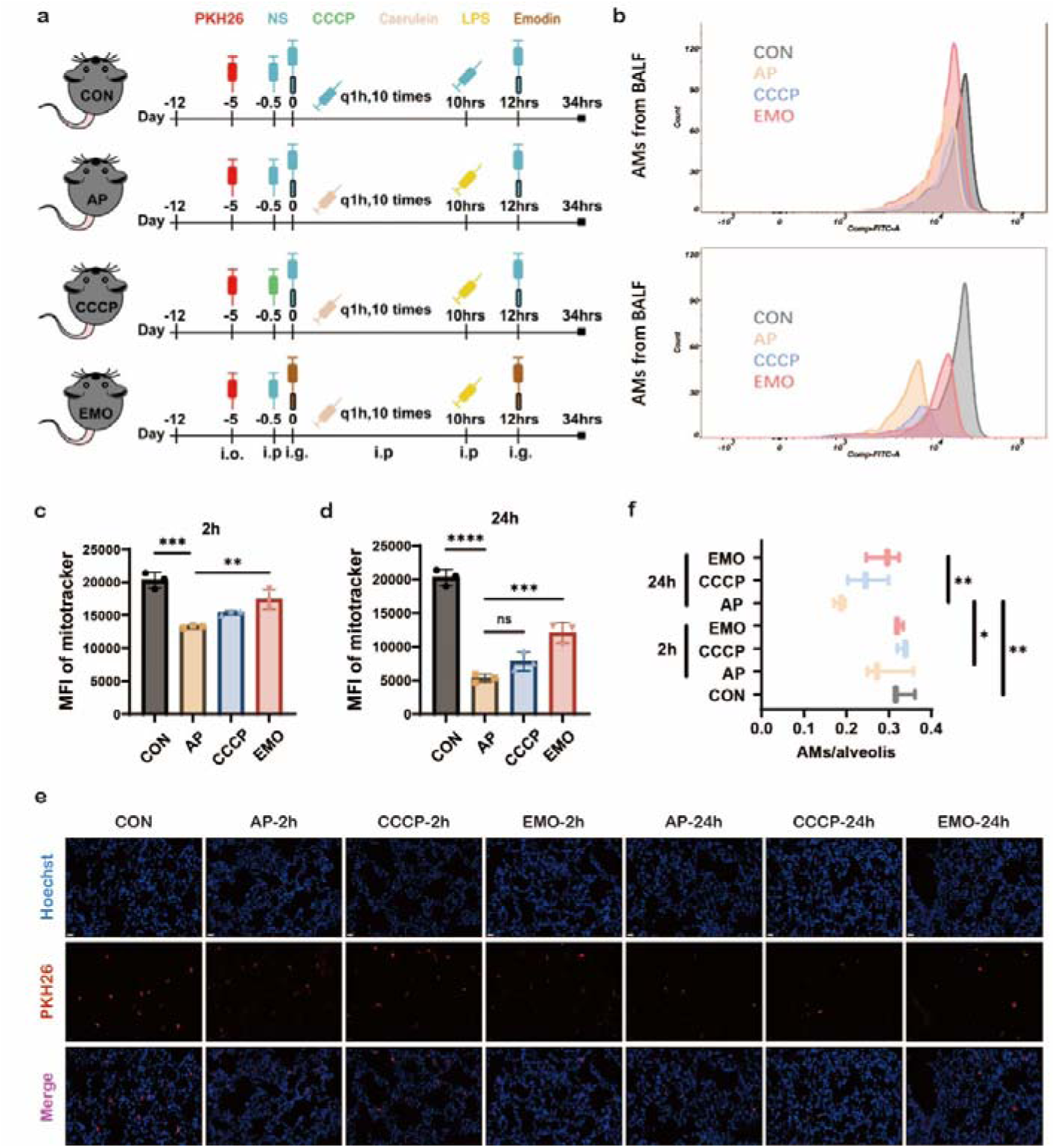
EMO impacted mitochondria changes in AMs during acute pancreatitis. (a) Schematic of the acute pancreatitis model and workflow. (b) Flow cytometry assessment of mitochondrial density in BALF in control or paralleled experiments at 2h and 24h after the model was established. (c) Quantification of mitotracker fluorescence in AMs at 2h after caerulein and LPS injection. (d) Quantification of mitotracker fluorescence in AMs at 24h after caerulein and LPS injection. (e) Representative images of frozen sections in lung from each group at the indicated time points. Hoechst (blue) marks the nucleus, PKH-AMs are shown in red. Empty (black) spaces are alveolar spaces. Scale bars: 20 μm. (f) Quantification of AMs/alveolis from each group. n=6/group.

**Figure 4.**
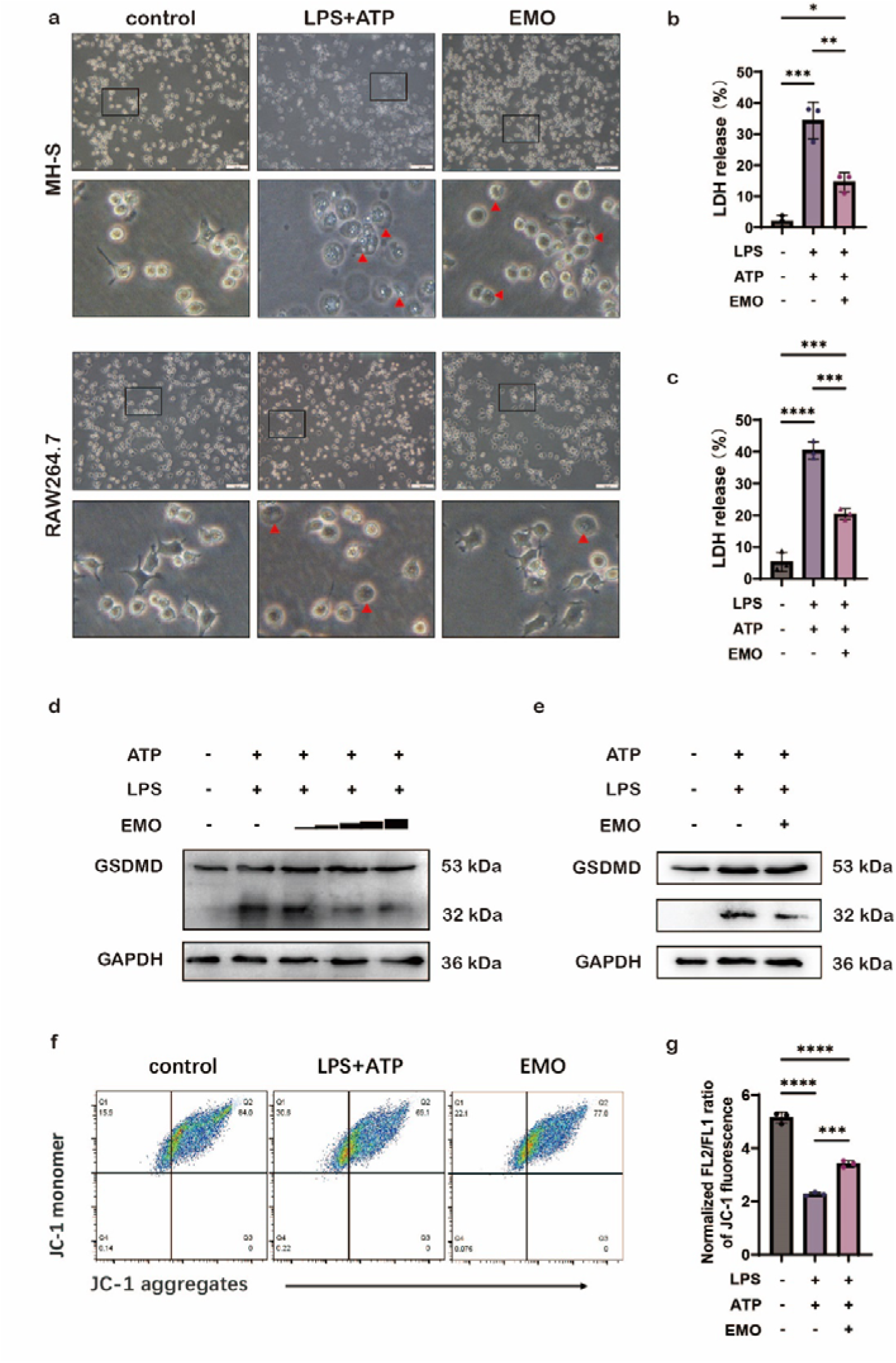
EMO inhibits pyroptosis in MH-S and RAW264.7, and changes the voltage of mitochondria in MH-S. (a) The morphology and number of pyroptotic MH-S and RAW264.7 were observed under a microscope. MH-S was pretreated with EMO (5μM, 2h), primed with LPS (500ng/ml, 4h), and followed by stimulated with ATP (2mM, 1h), while RAW 264.7 pretreated with EMO (10 μM, 2h), primed with LPS (500ng/ml, 12h) and followed by stimulated with ATP (2mM, 0.5h). Scale bars represent 100 μm. (b) Quantification of LDH in MH-S cultural supernatant. (c) Quantification of LDH in RAW264.7 cultural supernatant. (d) Immunoblots of GSDMD of MH-S protein lysates, when pretreated with various doses of EMO (5, 10, 15 μM, 2h), primed with LPS (500ng/ml, 4h) and followed by stimulated with ATP (2mM, 1h). n=3/condition. (e) Immunoblots of GSDMD of RAW264.7 protein lysates, when pretreated with various doses of EMO (10 μM, 2h), primed with LPS (500ng/ml, 12h) and followed by stimulated with ATP (2mM, 0.5h). n=3/condition. (f) Analysis of mitochondrial membrane potential levels in MH-S by JC-1 fluorescence. n=3/condition. (g) Quantification of mitochondrial membrane potential.

Emerging studies have reported that abnormality of mitochondrial membrane potential is tightly associated with the severity of inflammation in many inflammatory diseases. To investigate how EMO regulates mitochondrial voltage in macrophages, we utilized the JC-1 system to assess the changes of mitochondrial membrane potential after the stimulation of LPS plus ATP. Our results showed that LPS/ATP stimulation significantly decreased the mitochondrial membrane potential and reduced JC-1 aggregation in mitochondrial matrix (Fig. 4f-g). The existence of green fluorescence mon-omers in Fig. 4f indicated a mitochondrial membrane potential abnormality in LPS/ATP group. However, with 2h of EMO pretreatment, there is an increase in mitochondrial membrane potential evidenced as the gradual transition of fluorescence color to red (the alternative mitochondria-specific markers, MitoTracker Deep Red and Total MitoTracker Green, were used to stand for the differentiation of healthy mitochondrial). And the statistical analysis in Fig. 4g further confirmed the findings of the above.

### EMO inhibits NLRP3 inflammasome activation of macrophage in vitro

Then, we investigated the potential effects of EMO on NLRP3 inflammasome activation in macrophage. In Fig. 5a, pretreatment of EMO for 2h induced significant decreases of NLRP3, pro-Caspase-1, and Caspase-1 expression in a dose-dependent manner (2.5, 5, 7.5 μM) in MH-S macrophages in vitro. However, in RAW264.7 macrophages, there is a reduction of NLRP3 and Caspase-1, but no difference on pro-Caspase-1 levels (Fig. 5b). Furthermore, EMO dramatically reduced IL-1β secretion associated with LPS/ATP stimulation both in MH-S and RAW264.7 macrophages (Fig. 5c-d). In addition, based on dual-labeling of PI and Hoechst, EMO presents significant protective effects on cell death both in MH-S and RAW264.7 in the stimulation of LPS/ATP (Fig. 5e-h). Our results indicated the protective role of EMO on LPS/ATP induced cell death in murine macrophages, might function through inhibiting the activation of the NLRP3 inflammasome.

**Figure 5.**
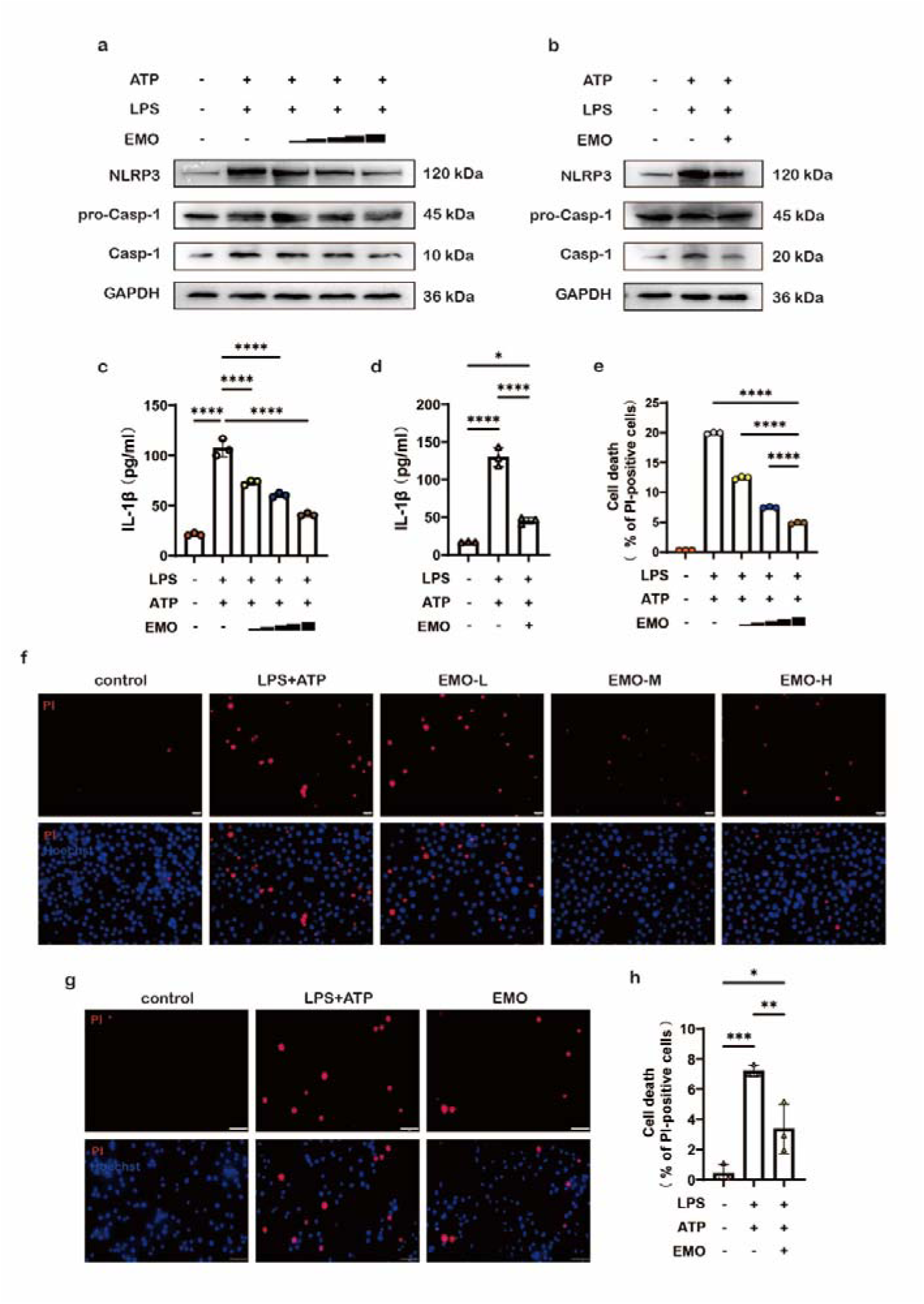
EMO suppresses the activation of NLRP3 inflammasome and prevents cell death in MH-S and RAW264.7. (a) Immunoblots of NLRP3 and caspase-1 on protein lysates from MH-S pre-treated with various doses of EMO (2.5, 5, 7.5 μM, 2h), primed with LPS (500ng/ml, 4h) and followed by stimulated with ATP (2mM, 1h). n=3/condition. (b) Immunoblots of NLRP3 and caspa-se-1 on protein lysates from RAW 264.7 pretreated with various doses of EMO (10 μM, 2h), primed with LPS (500ng/ml, 12h) and followed by stimulated with ATP (2mM, 0.5h). n=3/ condition. (c) Quantification of IL-1β in cultural supernatant by ELISA. (d) Quantification of IL-1β in cultural supernatant by ELISA. (e) Quantification of PI-positive cells. n=3/condition. (f) The cell death immunofluorescence photographs were produced by staining with Hoechst 33342 (5 μg/ml, blue) and PI (2 μg/ml, red). Scale bars: 20 μm. (g) Representative immunofluorescence images of cell death. Scale bars represent 50 μm. (h) Quantification of PI-positive cells. n=3/group.

### EMO enhanced mitophagy which leads to an alleviation of mitochondrial damage in macrophage in vitro

To explore EMO therapeutics on mitochondrial damage, we utilized dual labeling of MitoTracker Green positive and MitoTracker Deep Red negative staining which represented the percentage of damaged mitochondria. The data in Fig. 6a-b showed that LPS/ATP stimulation significantly increased mitochondrial membrane potential to 18% compared to 3.46% in control group. However, 5μM of EMO pretreatment for 2h, dramatically decrease the mitochondrial membrane potential to 14,7% in MH-S cells. The results were also validated in RAW264.7 macrophages (Figs. 3a and b). On the other hand, the combination of LPS/ATP stimulation induced an increase of reactive oxygen species (ROS) resulted from destroyed mitochondria (Fig. 6c). EMO pretreatment could significantly reduce the ROS level based on the quantification and flow cytometry analysis of MitoSOX in MH-S macrophages (Fig. 6c-d). In addition, we also assessed the ratio of mitochondrial DNA (mtDNA) and nuclear DNA (nDNA) to investigate mitochondrial damage induced by LPS/ATP or CCCP stimulation, and the data in Fig. 6e-f suggested a significant protects of EMO on increasing the mtDNA/nDNA ratio in MH-S macrophages, sequentially avoid mitochondrial damage. Mitophagy, a subtype of autophagy, plays a vital role in maintaining the balance of reactive oxygen species in mitochondria of macrophage. In the current study, we measured one of the mitophagy indicator-LC3II/LC3I, and EMO pretreatment showed a dramatic effect on increasing LC3II/LC3I levels which indicated a protective therapeutic of EMO in LPS/ATP associated mitophagy disorder (Fig. 6g). Moreover, mitochondria morphology (such as: cristae, telltale signs of double membrane autophagosomes) were also improved after EMO pretreatment (Fig. 6h). To further investigate the mitophagy alteration by EMO, another mitochondrial mark-er-TOMM20 was used in MH-S macrophage with LPS/ATP stimulation. In Fig. 6i, an increase of colocalization of TOMM20 with DRP1 (dynamin-related protein, a key mediator of mitochondrial fission) in MH-S macrophage indicated EMO exhibited consistent protects on mitochondria. Additionally, the results in RAW264.7 macro-phage, EMO presented similar effects on LC3 level and mitochondria marker tracking. Our data suggested that the protective effects of EMO in LPS/ATP stimulated in vitro macrophage model could function as inhibiting NLRP3 inflammasome and enhancing mitophagy, sequentially avoid further damage of mitochondria.

**Figure 6.**
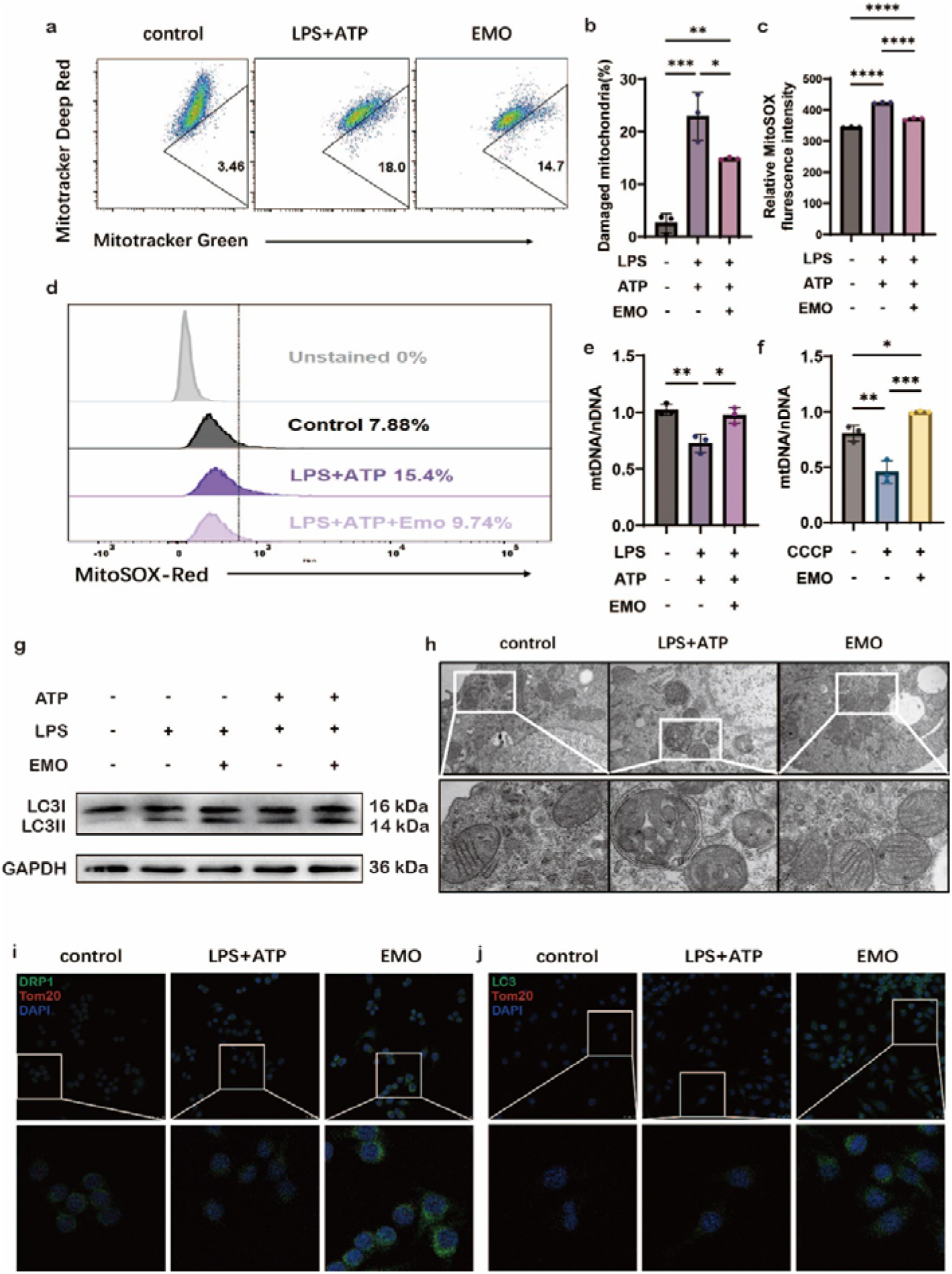
EMO induces mitophagy as a means to reduce mitochondrial injury. (a) MH-S mitochondrial membrane potential levels were analyzed using MitoTracker deep red and MitoTracker green staining. MH-S was pretreated with EMO (5μM, 2h), primed with LPS (500ng/ml, 4h), and followed by stimulated with ATP (2mM, 1h). n=3/condition. (b) Quantification of damaged mitochondria. (c) Quantification of MitoSOX. (d) Flow cytometry analysis of MitoSOX. n=3/condition. (e) The ratio of mitochondrial DNA (mtDNA) to nuclear DNA (nDNA) in MH-S was analyzed by qPCR to ascertain the mass of the mitochondria. n=3/condition. (f) The ratio of nDNA to mtDNA in MH-S was treated for 2 h with or without EMO (5uM) and CCCP (500nM). n=3/condition. (g) Immunoblots of LC3I and LC3II on protein lysates from MH-S pretreated with/without various doses of EMO (5μM, 2h), primed with LPS (500ng/ml, 4h) and followed by stimulated with/without ATP (2mM, 1h). n=3/condition. (h) The morphology of mitochondria in MH-S, both with and without EMO pretreatment and LPS/ATP treatment. n=3/condition. Scale bars: 0.5 μm. (i) The colocalization of DRP1 (green) and Tom20 (red) in MH-S. n=3/condition. Scale bars: 25 μm. (j) The colocalization of LC3 (green) and Tom20 (red) in MH-S. n=3/condition. Scale bars: 25 μm.

### Mitophagy inhibition reverses the protective effects of EMO against mitochondrial damage and inflammation in murine macrophages

To elucidate if EMO’s protects involved manipulating alveolar macrophage mitophagy during the AP-ALI, we utilized an autophagy/mitophagy inhibitor, 3-methyladenine (3-MA) in in vitro. Interestingly, with 3-MA treatment, there is a significant increase in ROS and mtDNA levels together with high accumulation of damaged mitochondria after LPS/ATP stimulation both in 3-MA group and EMO/3-MA group in MH-S macrophage (Fig. 7a-c). Furthermore, 3-MA treatment also blocked the alleviation of EMO-induced IL-1β release in the MH-S (Figs. 4). Specifically, 3-MA treatment showed significant inhibition on EMO-induced increase of LC3II and LC3I expression in both MH-S and RAW264.7 cells, which suggested that EMO-associated protective effects in macrophage functioned as manipulating mitophagy pathway (Fig. 7d-e). Moreover, mitophagy inhibition by 3-MA also presented considerable impacts on EMO induced IL-1 beta, TNF alpha, and IL-6 expression, but no influence on IL-18 level (Fig. 7i). Notably, the observation of the colocalization of Mitotracker and lysotracker indicated that EMO could increase mitophagy activity and alleviate mitochondrial damage; however, 3-MA treatment showed a significant decrease of Mitotracker signals and an increase of lysotracker amount in Fig. 7j. Our data here suggested that EMO’s protectives against mitochondria damage in macrophage are through regulating the NLRP3 inflammasome-mitophagy pathway.

**Figure 7.**
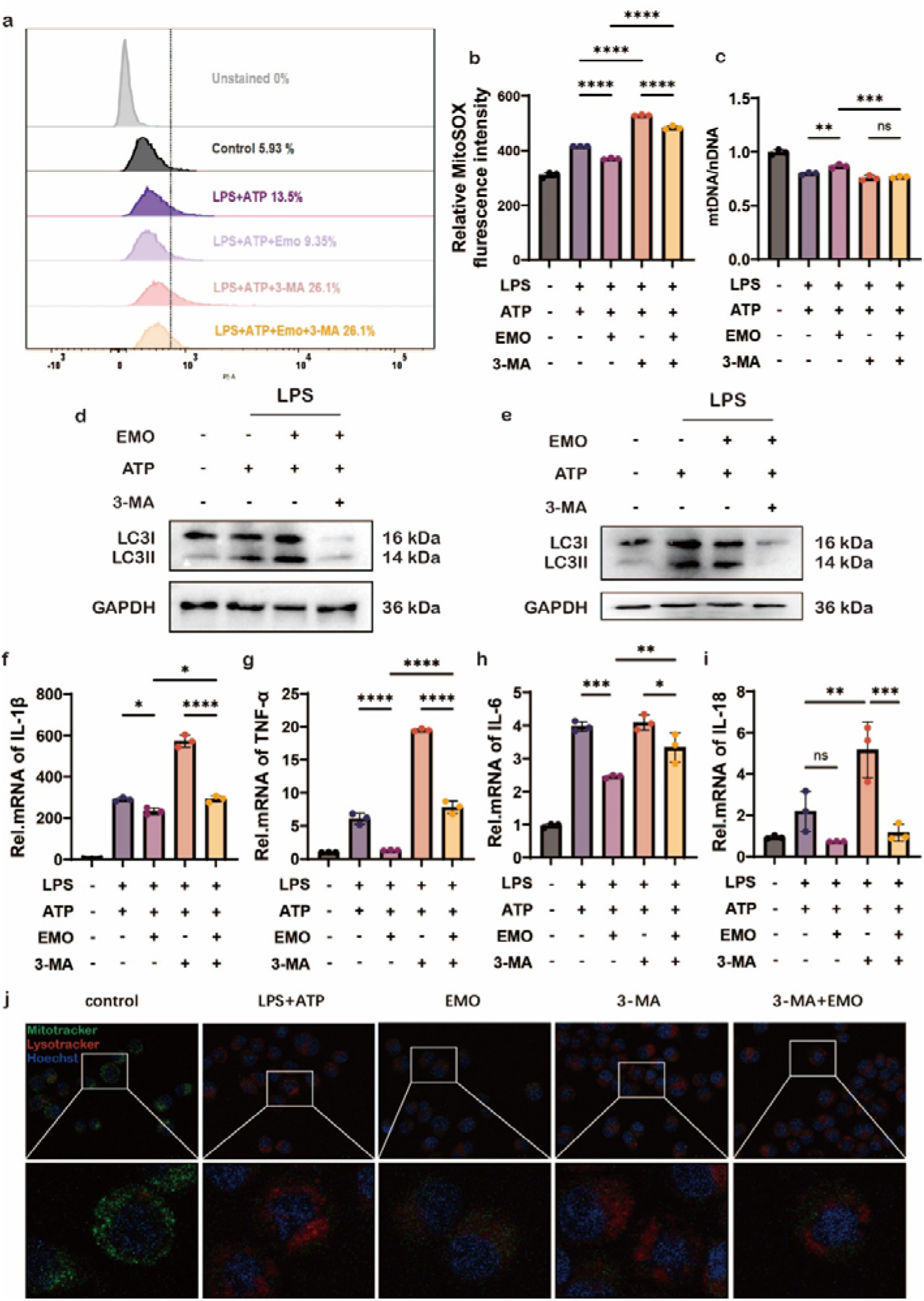
Inhibiting autophagy in murine macrophages counteracts EMO’s protective effects against inflammation and mitochondrial damage. (a) Analysis of MitoSOX levels in MH-S cells were treated with 3-MA (5mM, 1h) before EMO (5μM, 2h) treatment, LPS (500 ng/ml, 4h) and ATP (2mM, 0.5h) treatment. (b) Quantification of relative MitoSOX fluorescence intensity. (c) The ratio of mtDNA to nDNA in MH-S was analyzed by qPCR to ascertain the mass of the mitochondria. (d) Western blot analysis of LC3I/II protein expression in MH-S cells. n=3/condition. (e) Western blot analysis of LC3I/II levels in RAW264.7 cells, when treated with 3-MA (5mM, 1h) before EMO (5μM, 2h) treatment, LPS (500 ng/ml, 4h) and ATP (2mM, 0.5h) treatment. n=3/condition. (f-i) qPCR analysis of IL-1 beta, TNF alpha, IL-6, and IL-18 expression in MH-S. (i) The colocalization of Mi-totracker (green) and Lysotracker (red) in MH-S. n=3/condition. Scale bars represent 25 μm.

## Discussion

Till now, the pathophysiological mechanism of systemic injury in AP is still poorly understood. It has been well documented that lung is the most frequently involved organ associated with AP induced systemic damage^16^. Emerging studies showed that profiles of monocytes and macrophages are differentially altered during pancreatitis pathogenesis^17,18^. AMs monitor the luminal surface of the epithelium, together with epithelial cells, contributing to set the threshold and the quality of the innate immune response in the lung mucosa^19^. On the other hand, overactivation of AMs could also induce certain tissue damage, therefore it is quite necessary to elucidate how exactly AMs are involved in AP-ALI. With the unique phagocytic characteristic of macrophages, it was marked with PKH26 fluorescence in vivo prior to being observed in frozen slices. Benefited with this technique, we could not only track the macrophage location, but also measure the number of labeled macrophages in situ compared with the amount of macrophages in alveolar space.

Previous studies found that alveolar macrophages efficiently intercept particles, bacteria, and foreign materials, contributing to alveolar defense and clearance. Nevertheless, under the condition of a decrease of AMs-to-alveoli numbers ratio, the “self-cleaning” ability of AMs would dramatically decrease in inflammatory diseases, such as: acute pancreatitis, etc. Specifically in the initial stage of AP-ALI, once the alveolar equilibrium was disturbed, molecules from the collectin family (such as: SP-A, SP-D, and C1q) would facilitate phagocytosis and inflammation by binding the apoptotic cells via calreticulin-CD91 in AMs. Notably, there exists a critical time range for the contacts of alveolar epithelium with deceased cells, bacteria, fungi, and particles. And the process of those contacts will induce the release of multiple chemokines and neutrophils recruitment. In the present study, only a single AM was detected in approximately every three alveoli in the AP model which indicated a remarkable decrease across the time even though less impact on the resident AMs in the early stage. Based on these results, we hypothesized if the number of resident AMs could be chosen as a potential marker for the severity of AP-ALI. Using flow cytometry technique, we assessed mitochondria levels in resident AMs. The results showed that a decrease of mitochondria level was observed at 2h after caerulein-induced acute pancreatitis mouse model, and a continuous reduction till 24h after the event. It has been demonstrated that mitochondrial plays a critical role in regulating inflammatory responses through its genuine DAMPs which serve as physical scaffold for activating pattern recognition receptors (PRRs)^9^. Specifically, mitochondria provide a distinctive framework for DAMP redistribution, PRR signaling, and inflammation responded to cellular stress; on the other hand, these processes could stimulate innate and adaptive immune reactions to maintain the homeostasis^10^. However, during the inflammatory cascading amplification of AP, quite a few surface receptors have been dramatically increased. Meanwhile, multi-cellular responses (such as: autophagy, caspase) were also activated in response to RCD-associated mitochondrial outer membrane permeabilization (MOMP) to maintain homeostasis of AMs. Further comparison from our results indicated that alteration of mitochondrial flux initiated earlier than those changes in resident AMs amount in AP-ALI. Therefore, we performed an in vitro study to identify the correlation between pyroptosis and mitophagy in AP-ALI mouse model. The data in the current study evidenced that mitochondrial levels in AMs might be used as an index for acute lung injury.

Our previous studies have demonstrated that EMO impacted the progress of AP-ALI through its anti-inflammatory characteristic. In the current study, we investigated how EMO exerts its protective therapeutics on mitochondria homeostasis in AMs after AP-ALI. To do so, we constructed in vitro macrophage cell models treated with LPS/ATP. The results showed that pretreatment of EMO significantly decreased the accumulation of uncleared damaged mitochondria and subsequentially inhibited pyroptosis. Furthermore, EMO pretreatment also enhanced mitophagy in macrophages, which should induce several benefits as follows: first, the greater number of healthy mitochondria, the better for the energy requests of activated immune cells. The consequence should support the switch of cell metabolism from a resting state to high metabolic state and maintain energy homeostasis; then maintaining normal permeability of mitochondrial membrane by EMO supported the characteristics of an-ti-inflammation in macrophages. In addition, EMO reduced the accumulation of ROS and mtDNA in the cytoplasm (mtDAMPs), and further inhibited NLRP3 inflammasome formation. Other reports also mentioned that EMO could activate Par-kin-mediated mitophagy and lead to delay heart aging^20^. Collectively, our data elucidated the therapeutic effects EMO on alleviating AP-ALI function as promoting mitophagy in AMs (Fig. 8), which provided a novel therapeutic target and extended EMO’s beneficial effects during the process of inflammatory diseases.

**Figure 8.**
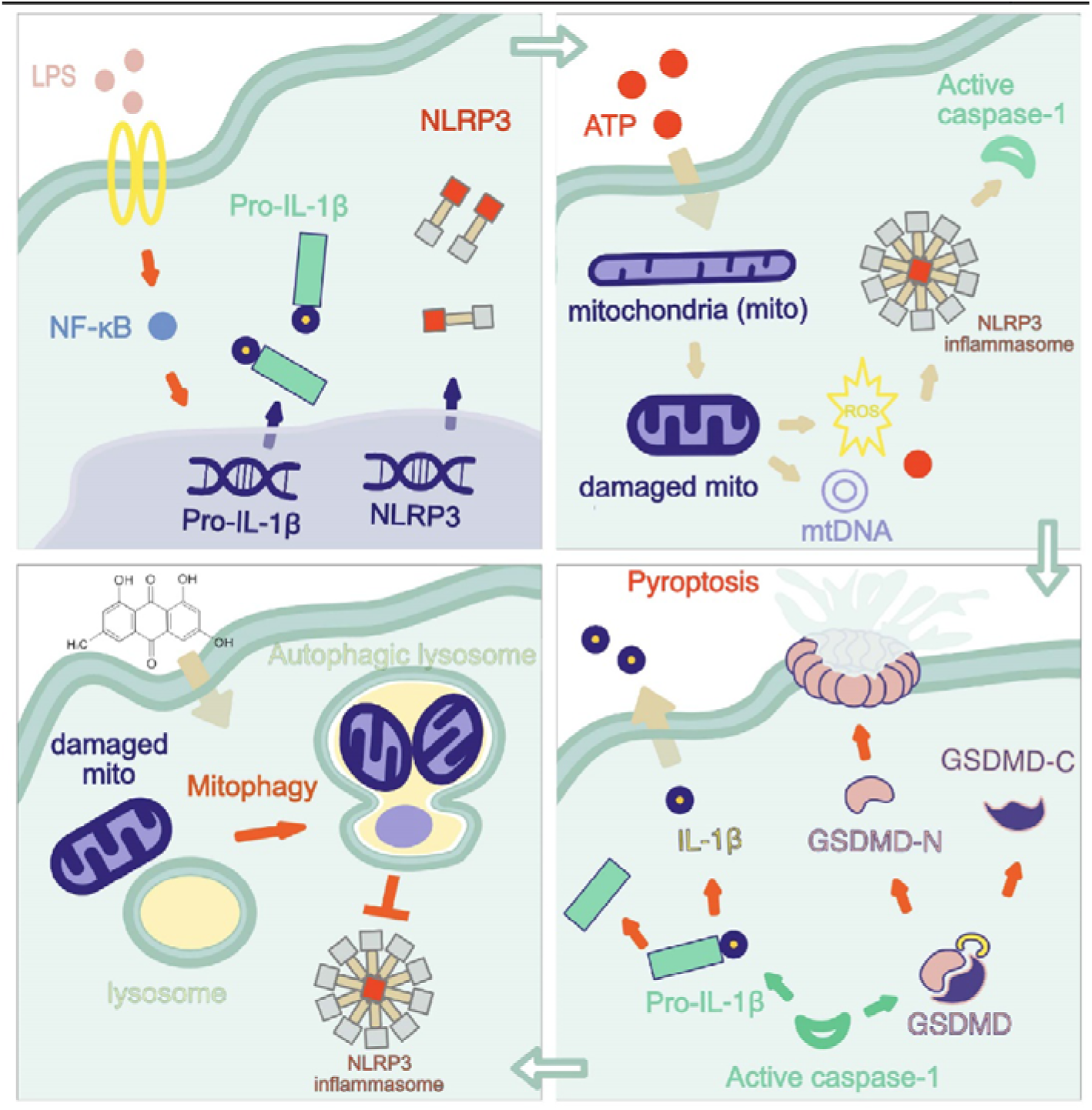
The regulatory mechanisms of EMO on alveolar macrophage in AP-ALI.

## Materials and Methods

### Animals

C57BL/6 male mice (12 weeks) were fed with a standard chow diet and housed in a temperature- and humidity-controlled facility. The animal experiments were conducted using experimental protocol AEE19003, which was authorized by the Research and Animal Ethics Committee of Dalian Medical University. The experiments were carried out in agreement with national and international criteria for the care and use of laboratory animals.

### AP-ALI mouse model

Acute pancreatitis was induced by intraperitoneal injection of caerulein (100 μg/kg, i.p. every hour ×10) which has been extensively documented^21^, and 10mg/kg LPS (E. coli serotype O55:B5) was intraperitoneal injected right after the last injection of caerulein. Control mice only received vehicles (5 ml/kg, 1×PBS). EMO (40 mg/kg body weight) was administered twice (right before the first dose of caerulein, and two hours after the last dose of caerulein) through gastric gavage. Samples included blood, pancreas, lung, and bronchial alveolar lavage fluid were collected at 2h and 24h postinjection with LPS.

### AMs in vivo labeling technique

PKH26 phagocytic cell labeling followed the manufacturer’s instructions. Briefly, a 100 μM solution of PKH26 was diluted to a concentration of 0.5 μM using diluent B. Mice were subjected to anesthesia using 3% isoflurane for 3 min. Subsequently, the mice were promptly hung by their incisors at a 60° angle, and 75 μL of a 0.5 μM PKH26 solution was accurately transferred into the posterior part of the throat using a pipette. Then, observe the movement and behavior of animals inside the cages until walk normally.

### Cell culture

MH-S cells (MH-S is a macrophage, alveolar cell line that was isolated from the lung of a 7-week-old, male mouse.) maintained in RPMI-1640 medium supplemented with 10% fetal bovine serum (FBS), and RAW264.7 cells (a macrophage cell line that was established from a tumor in a male mouse induced with the Abelson murine leukemia virus.) maintained in DMEM with 10% FBS, were cultured at 37LJ in a humidified 5% CO2 incubator.

### Isolation of AMs in murine bronchoalveolar lavage fluid (BALF)

Murine alveolar macrophages were harvested from BALF of C57BL/6 mice. Briefly, Mice were euthanized with CO2. The trachea was exposed and cannulated with a 24G catheter (Wego). 0.8 ml ice-cold PBS with 100 μM EDTA was gently infused followed by extraction. A total volume of ∼3.5-3.8 mL/mouse was collected. Then, BALF fluid was centrifuged at 1500RPM at 4LJ for 10 min. The supernatants were collected and assessed by spectrophotometer for some experiments turbidity. After being subjected to ACK lysis (BD) for 2 min, the cell pellets were washed in 2% FBS and stored on ice until staining. Protein in supernatants was quantified by using BCA Protein Assay Kit, and total cholesterol was quantified by using Mouse Cholesterol Kit following manufac-turer’s instructions.

### Flow cytometry assay

Cells isolated from BALF were stained with 200 nm MitoTracker™ Green FM for 20 min. After incubation with viability using viability dye, Cells were incubated with blocking antibody (2.4G2, BD) for 5 min, as per manufacturer’s instructions. Cells were then stained with specific antibody cocktails at 4LJ. Negative Control Compensation Particles Set was also performed. For MH-S and RAW264.7 cells, samples were assessed by the LSRFortessa cell analyzer (BD) and FlowJo software.

### Inflammasomes activity assessment

MH-S or RAW264.7 cells were seeded in 12-well plates with cell density of 1×105 cells/ml. MH-S cells were pretreated with EMO (5μM, 2h) combined with LPS (500ng/ml, 4h), sequentially stimulated with ATP (2mM, 1h); RAW 264.7 cells were pretreated with EMO (10 μM, 2h) primed with LPS (500ng/ml, 12h), and then stimulated with ATP (2mM, 1h). The inflammasome activity in the supernatants were collected 1 hour after the treatment and analyzed with enzyme-linked immunosorbent assay (ELISA) kits.

### Cell viability assay

The cell viability of MH-S and RAW264.7 was assessed using the CCK-8 kit. In short, MH-S and RAW264.7 cells were seeded in 96-well plates at ∼1×10^5^ cells/well. 12h later, cells were treated with indicated concentration of EMO for 6/12h, then incubated with 10 μl of CCK-8 solution at 37LJ for 30 min. At the indicated wavelength, the mixture’s absorbance was measured.

### Pyroptosis assay

PI incorporation was used to detect pyroptosis according to a previous documentary. The cells were stained with PI (2 μg/ml) and Hoechst 33342 (5 μg/ml) for 10 min after being treated with EMO, LPS, and ATP, as previously stated. Olympus IX73 was used to examine stained cells. Analysis was performed based on a minimum of six random fields. The LDH cytotoxicity test kit was used to determine the levels of LDH released in the culture supernatants.

### Transmission electron microscopy (TEM)

Cells were fixed at room temperature with 2.5% glutaraldehyde (Absin). Then, it was rinsed three times with PBS, and subjected to an escalating progressive sequence of ethanol treatments for 8 minutes. Samples were then infiltrated with resin and polymerized. The images were acquired with transmission electron microscopy.

### Enzyme Linked Immunosorbent Assay (ELISA)

The levels of IL-1β and TNF-α in either serum, BALF, or cultured supernatants were determined with ELISA kit following the manufacturer’s instructions.

### Western blot assay

Cells were lysate with RIPA buffer containing a protease inhibitor cocktail (NCM Bio-tech). Protein concentrations were measured using a BCA protein assay kit. After undergoing SDS-PAGE electrophoresis, the protein samples were transferred to nitro-cellulose membranes and blotted in 5% non-fat milk for an hour at room temperature. After incubating with the appropriate HRP-conjugated antibodies for an hour at room temperature, protein signals were observed via chemiluminescence detection, and the intensity was analyzed by ImageJ software.

### Superoxide and mitochondrial membrane potential evaluation

Cells were treated with 3-MA (5 mM, 1h) and stimulated as previously stated. Mito-SOX red, which be used as a fluorescent indicator to specifically detect superoxide, producing red fluorescence and JC-1 fluorescent probe, an ideal fluorescent probe widely used to detect mitochondrial membrane potential, was assembled in accordance with each kit’s instructions, respectively. Samples were then analyzed by flow cytometry as described above.

### DNA and RNA isolation and quantitative RT-PCR

A mitochondrial D-loop-specific primer allowed us to determine the mtDNA/nDNA ratio. ONE-4-ALL Genomic DNA Mini-Preps Kit was used to purify MH-S cellular DNA according to the manufacturer’s instructions. Total RNA was isolated by Trizol reagent and reverse transcribed DNA was amplified by Monscript RTIII All-in-One Mix with dsDNase. Quantitative RT-PCR was performed with RapidStart Universal SYBR Green qPCR Mix using a LineGene 9600 Plus fluorescence quantitative thermal cycler (Bioer).

### Immunofluorescent staining

Cells were fixed with 4% paraformaldehyde at 4LJ for 10 min and permeabilized by PBS (0.1% Triton-100). Then, blocked by PBST (PBS plus 0.1% Tween-20) at 4LJ for 30 min following the incubation with specific primary at 4LJ overnight. Next, samples were incubated with appropriate second fluorescent antibodies (Alexa-488, Alexa-568) and DAPI. The MitoTracker Green FM (200 nM) and Lyso-Tracker Red (50 nM) were performed according to the manufacturer’s instructions. All images were performed by using an Olympus BX63 confocal microscope system.

### Wet/dry eight (W/D) ratio calculation

Briefly, the upper part from right side of lung was removed and weighed immediately for wet weight measurement. Then, tissue was placed in an oven at 70°C for 24h to completely dry the moisture. The dry weight of lung tissue was evaluated, and W/D ratio was obtained.

### Statistical analysis

The PRISM program was used to do statistical analysis. Unless specified otherwise in the image caption, the mean standard deviation represents all results from in vitro and in vivo studies. An unpaired Student’s t-test was used to analyze differences between the two groups, and a one-way ANOVA was conducted for groups with multiple variables and more than two variables. At * P<0.05, ** P<0.01, *** P<0.001, **** P<0.0001, differences were deemed statistically significant.

## Conclusions

The current study indicated that 1) mitochondria status determined the fate of resident AMs and is critical in AP-ALI in mice; 2) monitoring and proactive intervention on mitophagy could protect mitochondria damage and sequentially improved AP; 3) EMO protective therapeutics in AP-ALI function as inhibiting NLRP3 inflammasome activation, increasing mitophagy activity, eliminating ROS, and impacting pyroptosis process. Our results suggested that EMO could be used as a reliable anti-inflammatory and mitophagy enhancer during the AP-ALI, which should shed light on therapeutic improvements for inflammatory diseases in clinics.

## Acknowledgments

The authors thank Dr. Huiyi Song and Dr. Fangyue Guo for their significant contributions to the technical support of this paper.

## Author contributions

ZC and HLC conceived and designed the experiments. ZC, XCD, YWS, BWL, YLL and HYW performed experiments. ZC and XCD analyzed the data. HLC secured funding. ZC and XCD wrote the paper. All authors have read and approved the final manuscript.

## Data availability

The datasets used and/or analyzed during the current study are available from the corresponding author upon reasonable request.

## Additional information Competing interest

The authors declare no competing interest.

## Supplementary Materials

Figs. 1. Pancreatic histopathological injury, score, lipase level and flow cytometry analysis. Figs. 2. Effect of EMO on cell activity of MH-S and RAW264.7 cells. Figs. 3. Percentage of mitochondrial damage. Figs. 4. Concentration of IL1β. Supplementary Table S1. Information on antibodies, chemicals, peptides, critical commercial assays, experimental models, oligonucleotides, and software.

## Approval for animal experiments

This study was performed in line with the principles of the Declaration of Helsinki. Animal experiments were performed following experimental protocol AEE19003, approved by the Research and Animal Ethics Committee of Dalian Medical University (Dalian, China).

## References

1. Aldiabat, M., Kilani, Y., Arshad, I., Rana, T., Aleyadeh, W., Al Ta’ani, O., Aljabiri, Y., Alsakarneh, S., Abdelfattah, T., Alhuneafat, L., et al. (2023). Determinants and outcomes of acute pancreatitis in patients hospitalized for COVID-19: Early pandemic experience. Pancreatol. Off. J. Int. Assoc. Pancreatol. IAP Al 23, 926–934. 10.1016/j.pan.2023.10.012.

2. Lee, P.J., Lahooti, A., Culp, S., Boutsicaris, A., Holovach, P., Wozniak, K., Lahooti, I., Paragomi, P., Hinton, A., Pothoulakis, I., et al. (2023). Obesity and alcoholic etiology as risk factors for multisystem organ failure in acute pancreatitis: Multinational study. United Eur. Gastroenterol. J. 11, 383–391. 10.1002/ueg2.12390.

3. Szatmary, P., Grammatikopoulos, T., Cai, W., Huang, W., Mukherjee, R., Halloran, C., Beyer, G., and Sutton, R. (2022). Acute Pancreatitis: Diagnosis and Treatment. Drugs 82, 1251–1276. 10.1007/s40265-022-01766-4.

4. Kang, H., Yang, Y., Zhu, L., Zhao, X., Li, J., Tang, W., and Wan, M. (2022). Role of neutrophil extracellular traps in inflammatory evolution in severe acute pancreatitis. Chin. Med. J. (Engl.) 135, 2773–2784. 10.1097/CM9.0000000000002359.

5. Ge, P., Luo, Y., Okoye, C.S., Chen, H., Liu, J., Zhang, G., Xu, C., and Chen, H. (2020). Intestinal barrier damage, systemic inflammatory response syndrome, and acute lung injury: A troublesome trio for acute pancreatitis. Biomed. Pharmacother. Biomedecine Pharmacother. 132, 110770. 10.1016/j.biopha.2020.110770.

6. Zou, S., Jie, H., Han, X., and Wang, J. (2023). The role of neutrophil extracellular traps in sepsis and sepsis-related acute lung injury. Int. Immunopharmacol. 124, 110436. 10.1016/j.intimp.2023.110436.

7. Picca, A., Faitg, J., Auwerx, J., Ferrucci, L., and D’Amico, D. (2023). Mitophagy in human health, ageing and disease. Nat. Metab. 5, 2047–2061. 10.1038/s42255-023-00930-8.

8. Mohsin, M., Tabassum, G., Ahmad, S., Ali, S., and Ali Syed, M. (2021). The role of mitophagy in pulmonary sepsis. Mitochondrion 59, 63–75. 10.1016/j.mito.2021.04.009.

9. Zimmermann, A., Madeo, F., Diwan, A., Sadoshima, J., Sedej, S., Kroemer, G., and Abdellatif, M. (2023). Metabolic control of mitophagy. Eur. J. Clin. Invest., e14138. 10.1111/eci.14138.

10. Lyu, Y., Wang, T., Huang, S., and Zhang, Z. (2023). Mitochondrial Damage-Associated Molecular Patterns and Metabolism in the Regulation of Innate Immunity. J. Innate Immun. 15, 665–679. 10.1159/000533602.

11. Malainou, C., Abdin, S.M., Lachmann, N., Matt, U., and Herold, S. (2023). Alveolar macrophages in tissue homeostasis, inflammation, and infection: evolving concepts of therapeutic targeting. J. Clin. Invest. 133, e170501. 10.1172/JCI170501.

12. Hu, Y., Yang, L., and Lai, Y. (2023). Recent findings regarding the synergistic effects of emodin and its analogs with other bioactive compounds: Insights into new mechanisms. Biomed. Pharmacother. 162, 114585. 10.1016/j.biopha.2023.114585.

13. The versatile emodin: A natural easily acquired anthraquinone possesses promising anticancer properties against a variety of cancers - PubMed https://pubmed.ncbi.nlm.nih.gov/35637953/.

14. Emodin Ameliorates Severe Acute Pancreatitis-Associated Acute Lung Injury in Rats by Modulating Exosome-Specific miRNA Expression Profiles - PubMed https://pubmed.ncbi.nlm.nih.gov/38026528/.

15. Xu, Q., Wang, M., Guo, H., Liu, H., Zhang, G., Xu, C., and Chen, H. (2021). Emodin Alleviates Severe Acute Pancreatitis-Associated Acute Lung Injury by Inhibiting the Cold-Inducible RNA-Binding Protein (CIRP)-Mediated Activation of the NLRP3/IL-1β/CXCL1 Signaling. Front. Pharmacol. 12, 655372. 10.3389/fphar.2021.655372.

16. Jia, Y.-C., Ding, Y.-X., Mei, W.-T., Wang, Y.-T., Zheng, Z., Qu, Y.-X., Liang, K., Li, J., Cao, F., and Li, F. (2021). Extracellular vesicles and pancreatitis: mechanisms, status and perspectives. Int. J. Biol. Sci. 17, 549–561. 10.7150/ijbs.54858.

17. Hu, F., Lou, N., Jiao, J., Guo, F., Xiang, H., and Shang, D. (2020). Macrophages in pancreatitis: Mechanisms and therapeutic potential. Biomed. Pharmacother. Biomedecine Pharmacother. 131, 110693. 10.1016/j.biopha.2020.110693.

18. Cruz, A.F., Rohban, R., and Esni, F. (2020). Macrophages in the pancreas: Villains by circumstances, not necessarily by actions. Immun. Inflamm. Dis. 8, 807–824. 10.1002/iid3.345.

19. Tao, H., Xu, Y., and Zhang, S. (2023). The Role of Macrophages and Alveolar Epithelial Cells in the Development of ARDS. Inflammation 46, 47–55. 10.1007/s10753-022-01726-w.

20. Li, H., Liu, X., Meng, Z., Wang, L., Zhao, L., Chen, H., Wang, Z., Cui, H., Tang, X., Li, X., et al. (2022). Kanglexin delays heart aging by promoting mitophagy. Acta Pharmacol. Sin. 43, 613–623. 10.1038/s41401-021-00686-5.

21. Gorelick, F.S., and Lerch, M.M. (2017). Do Animal Models of Acute Pancreatitis Reproduce Human Disease? Cell. Mol. Gastroenterol. Hepatol. 4, 251–262. 10.1016/j.jcmgh.2017.05.007.

